# Designing Interpretable Convolution-Based Hybrid Networks for Genomics

**DOI:** 10.1101/2021.07.13.452181

**Authors:** Rohan Ghotra, Nicholas Keone Lee, Rohit Tripathy, Peter K. Koo

**Author notes:** Correspondence to: Peter K. Koo, < >. ICML 2021 Workshop on Computational Biology. Do not distribute.

## Abstract

Hybrid networks that build upon convolutional layers with attention mechanisms have demonstrated improved performance relative to pure convolutional networks across many regulatory genome analysis tasks. Their inductive bias to learn long-range interactions provides an avenue to identify learned motif-motif interactions. For attention maps to be interpretable, the convolutional layer(s) must learn identifiable motifs. Here we systematically investigate the extent that architectural choices in convolution-based hybrid networks influence learned motif representations in first layer filters, as well as the reliability of their attribution maps generated by saliency analysis. We find that design principles previously identified in standard convolutional networks also generalize to hybrid networks. This work provides an avenue to narrow the spectrum of architectural choices when designing hybrid networks such that they are amenable to commonly used interpretability methods in genomics.

## 1. Introduction

Convolutional neural networks (CNNs) are gaining popularity for regulatory genomic prediction tasks. To gain insights into the sequence features that influence their predictions, it is common to interpret CNNs by visualizing first layer filters or employing attribution methods (Kelley et al., 2018; Avsec et al., 2021b; Maslova et al., 2020; Atak et al., 2021). In practice, the infinite spectrum of design choices often makes it challenging to identify a suitable model.

Choice of architecture significantly influences the efficacy of interpretability methods. For instance, spatial information modulated by max-pooling can control the extent that first layer filters learn motif representations (Koo & Eddy, 2019). Moreover, an exponential function applied only to the first layer while using standard activations, such as a rectified linear unit (ReLU), in deeper layers, leads to learning robust motif representations and results in more trustworthy attribution maps (Koo & Ploenzke, 2021).

Recently, several hybrid networks that build upon convolutional layers with architectures developed for natural language processing, including bidirectional long-short-term memory (BiLSTM) (Quang & Xie, 2016; Minnoye et al., 2020), multi-head attention (MHA) (Li et al., 2020; Ullah & Ben-Hur, 2021), and transformer encoders (Ji et al., 2020; Avsec et al., 2021a), have demonstrated improved performance relative to pure CNNs. BiLSTMs can, in principle, capture long-range motif interactions. However, extracting these interactions is not straightforward. Alternatively, MHA, a key component of transformers, can provide an “interpretable” attention map to reveal learned activity between pairs of convolutional filters (Ullah & Ben-Hur, 2021). However, such an attention map is only intrinsically interpretable if the convolutional layer(s) learn robust motif representations and are identifiable in the attention maps.

Here we perform a systematic study to investigate how design choices in hybrid networks impact learning motif representations in first layer filters, as well as their reliability with saliency analysis (Simonyan et al., 2013). We find models that employ 2 design choices developed for standard CNNs, specifically larger pool sizes or exponential activations, consistently leads to learning robust motifs across all hybrid models. This work provides an avenue to narrow the scope of architecture choices when designing hybrid networks that are amenable to commonly used interpretability methods.

## 2. Experimental overview

To explore the role that model architecture influences performance and interpretability in a quantitative manner, we perform a systematic study across two tasks. Tasks 1 and 2 aim to address the question, “What extent do first layer convolutional filters learn motif representations?” and “How do design choices in complex networks influence the efficacy of attribution methods?”, respectively.

### Models

To systematically compare different hybrid architectures, we constructed each model using 3 stages (Fig. 1). Stage 1 includes 3 baseline modules: a single convolutional layer followed by a large max-pooling of size 24 (C24), 2 convolutional layers each with small max-pooling sizes of 4 and 6 (C4C6), and a convolutional layer followed by a BiLSTM each with small max-pooling of 4 and 6 (C4). The output lengths are the same for each module and thus enable a fair comparison. We append C24 and C4C6 with either MHA (Att), BiLSTM and MHA (LSTM-Att), or BiLSTM and different numbers of transformer encoders (LSTM-TransN, where N=1, 2,and 4). C4 was appended with either MHA or transformer layers, excluding the additional BiLSTM. Stage 3 includes a dense hidden layer followed by an output layer. As a control, we created models that directly connect Stage 1 baselines to Stage 3 outputs, without any attention modules. Each convolution and dense layer includes batch normalization prior to activations (Ioffe & Szegedy, 2015). Dropout (Srivastava et al., 2014) is incorporated after each convolution, BiLSTM, and MHA layer with a rate of 0.1 and dense layers with 0.5, with the exception of the transformer module (dropout rate=0.1).

**Figure 1.**
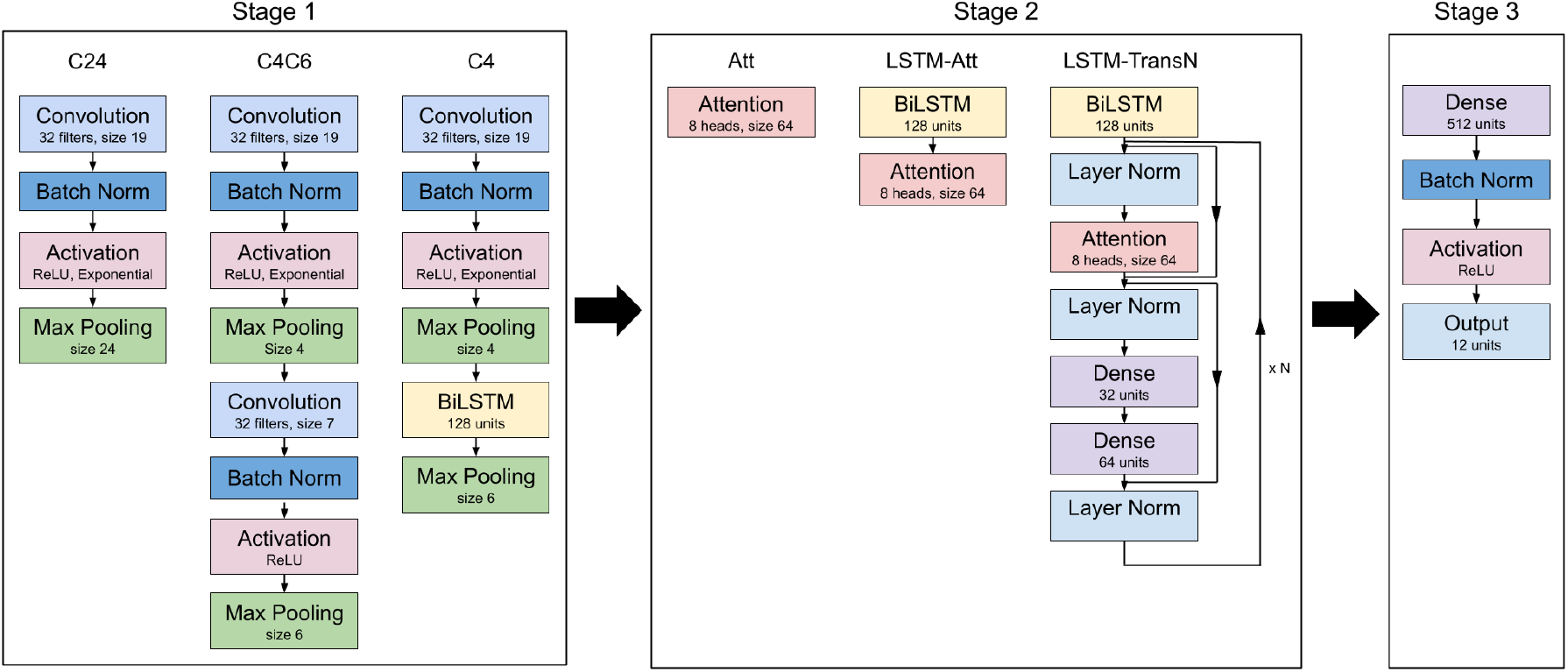
Model architectures. Each model employs a baseline module from Stage 1 and pairs it with an attention-based module from Stage 2, followed by the output module (Stage 3). The inputs to the model are one-hot encoded sequences of length 200.

We uniformly trained each model by minimizing the binary cross-entropy loss with mini-batch stochastic gradient descent (100 sequences) for 100 epochs with Adam updates using default parameters (Kingma & Ba, 2014). We decayed the learning rate by a factor 0.2 when the performance metric (AUPR for Task 1; AUROC for Task 2) did not improve for 4 epochs. Each model was trained 10 times with different random initializations according to (He et al., 2015). All results are taken from the top 3 models, based on best classification performance. This was necessary to remove runs where the deeper transformer models would simply not train due to poor initialization. Code availability: http://github.com/p-koo/hybrid_genomics.

### Task 1

We analyzed synthetic data that recapitulates a mutli-task classification of identifying simple TF binding sites from (Koo & Eddy, 2019). Briefly, 25,000 random DNA sequences, each 200 nucleotides long, were embedded with 1 to 5 binding sites, selected from a pool of 12 known motifs. The data was randomly split into training, validation and test sets with a 0.7, 0.1, and 0.2 split, respectively. After training, we visualized 1st layer convolutional filters via an activation-based alignment according to (Koo & Ploenzke, 2020). We quantified the extent that each filter matches a ground truth motif, i.e. true positive rate (TPR), using TomTom (Gupta et al., 2007). We also quantified how many filters that do not match any ground truth motifs have a statistically significant match to another motif in JASPAR (Mathelier et al., 2016), i.e. false positive rate (FPR).

### Task 2

We analyzed synthetic data that recapitulates a simple billboard model of gene regulation from (Koo & Ploenzke, 2021). Positive class sequences were embedded with 3 to 5 “core motifs” randomly selected with replacement from a pool of 5 known TF motifs. Negative class sequences were generated in a similar way but with the exception that the pool of motifs also includes 100 non-overlapping “background motifs” from JASPAR. Background sequences can thus contain core motifs; however, it is statistically unlikely for these sequences to resemble a positive class. 20,000 sequences were randomly split into train (0.7), valid (0.1), and test (0.2) sets. For this task, models were scaled down to 24 convolutional filters, BiLSTM and Attention size of 48, and dense layer of 96. After training, we computed a saliency map (Simonyan et al., 2013) for each positive-label sequence and multiplied it by the inputs, i.e. grad-times-input. We generated the distribution of saliency scores at positions where ground truth motifs were embedded and the distribution of saliency scores at other positions, as described previously (Koo & Ploenzke, 2021). We quantified the separation of these two distributions using 3 summary statistics: area under the receiver operating characteristic curve (AUROC), area under the precision-recall curve (AUPR), and signal-to-noise ratio (SNR). SNR is calculated by dividing the average saliency score at ground truth positions with the average across the 20 worst saliency scores at other positions for each sequence.

## 3. Results

We compared different hybrid networks, specifically combinations of 3 baselines and 3 attention-based modules (Fig. 1), to test the extent that architectural choices influence the ability to learn motif representations in first layer filters (Task 1) and provide reliable saliency maps (Task 2). Since we have ground truth, we can quantitatively measure the efficacy of the interpretability methods for each task.

### Task 1

We trained different hybrid networks and compared their classification performance on test data. All models tested in Task 1 achieved similar AUPR, with the exception of baselines that employ 4 transformer layers (Fig. 2). Due to the difficulty of training very deep networks, which are sensitive to poor intialization (Pennington et al., 2017), we only present results across 3 models that yield the highest prediction performance across 10 trials. We also quantified the TPR between filters and ground truth motifs as well as FPR to other motifs (see Section 2). Strikingly, we observed a significant improvement for C4C6- and C4-based models with exponential activations; the proportion of filters that match to ground truth motifs was significantly higher when exponential activations were used, compared to ReLU. On the other hand, C24-based models were able to learn motifs well for both activations, with a slight improvement for ReLU activations. Nevertheless, the FPR is largely the same, which suggests that many of the filters with exponential activations either learn the right motifs or none at all (i.e. no tomtom hits), creating sparsity, which is arguably a desirable property for model interpretability. These results are in agreement with a design principle identified previously with convolutional networks (Koo & Eddy, 2019) – models that employ small max-pooling can assemble whole motifs in deeper layers by spatially ordering partial motifs learned in the first layer; whereas, models that employ large max-pooling lose valuable spatial information. Thus, deeper layers cannot combine partial motifs; the only way it can reduce its loss is by learning complete motif representations. This demonstrates that although BiLSTM and MHA can capture long-range motif interactions, it only does so if the convolutional layer(s) learn robust motif representations; otherwise, it helps to assemble whole motifs from partial motifs. Evidently, the issues with small pooling can be overcome with exponential activations in first layer filters. Together, this demonstrates that design principles for CNNs to learn robust and identifiable motif representations in first layer filters can also be extended to hybrid networks.

**Figure 2.**
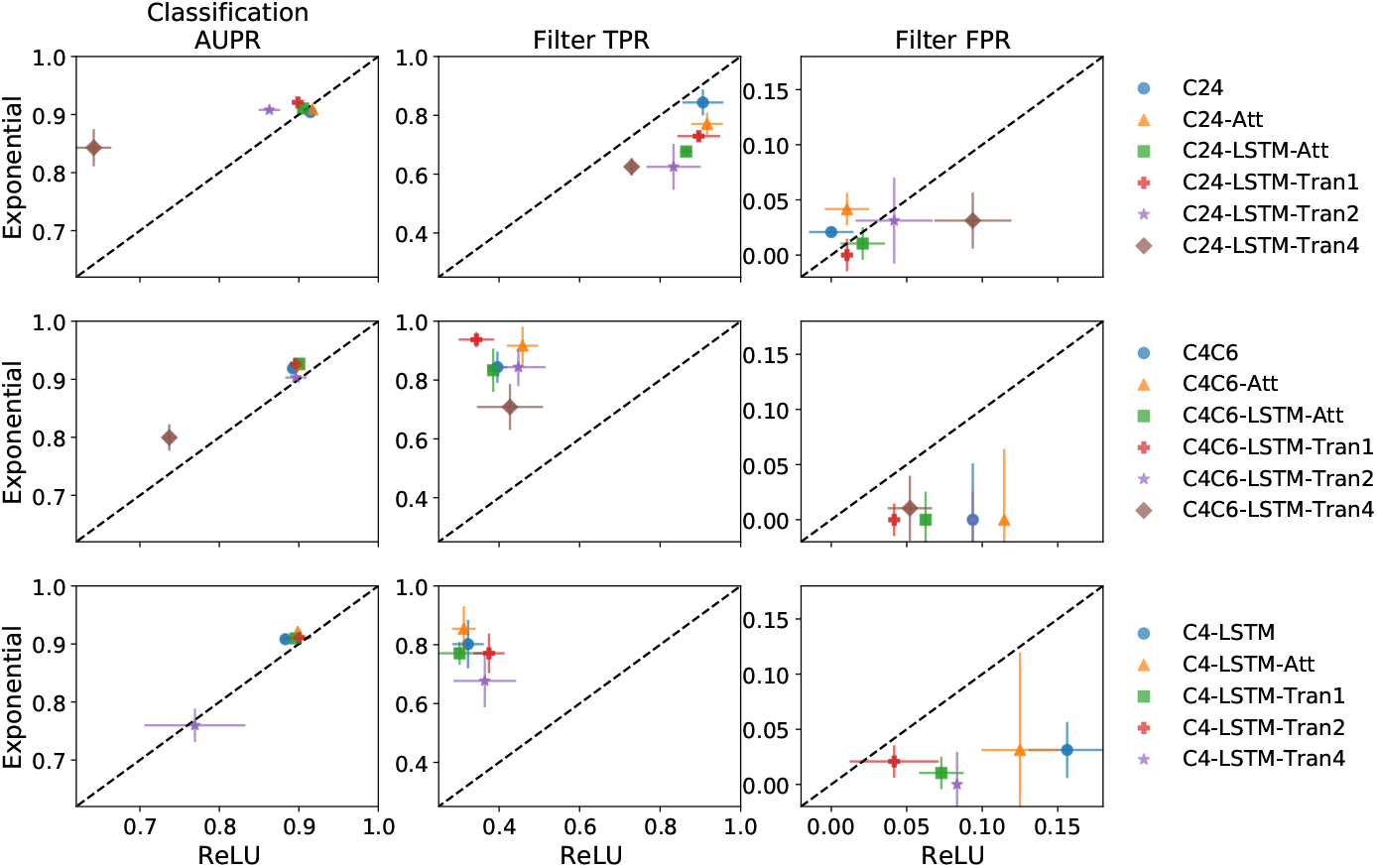
Task 1 filter analysis. Scatter plots of the classification AUPR on test data (left column), the true positive rate that first layer filters have a statistically significant match to a ground truth motif (middle column), and the false positive rate that first layer filters have a statistically significant match to a JASPAR motif but not any ground truth motif (right column) for hybrid networks with exponential activations versus ReLU activations. Each baseline architecture is shown on a different row. Error bars represent the standard deviation of the mean across 3 random intializations.

### Task 2

To explore the impact of architecture choice on the quality of saliency maps, we trained similar hybrid networks on a synthetic regulatory code dataset (Section 2). Evidently, models that employ small max-pooling (eg. C4C6 and C4) and exponential activations yield a significant improvement in both classification and interpretability performance compared to ReLU activations. On the other hand, only a slight gain was observed for C24-based models (Fig. 3). Unlike C4C6 and C4, C24-based models with ReLU activations learn full-length motifs in first layer filters; while exponential activations are needed to encourage C4C6 and C4-based models to learn better motif representations. Thus, our results suggest that models that learn robust motif representations (in the first layer) leads to improved reliability with attribution methods, in this case saliency maps, which is in agreement with previous results (Koo & Ploenzke, 2021).

**Figure 3.**
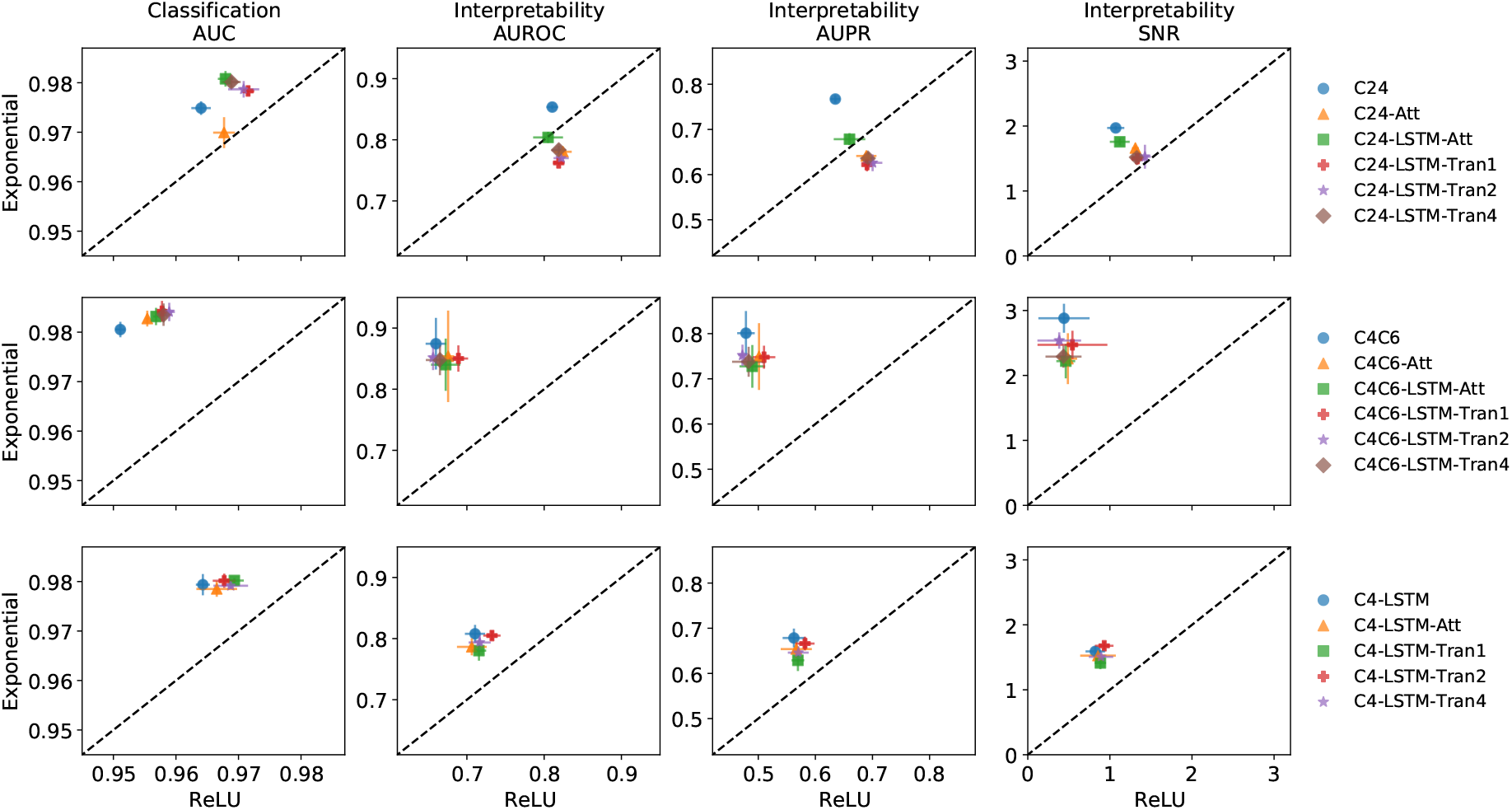
Task 2 saliency analysis. Scatter plots of the classification AUC on test data, the interpretability AUROC, AUPR, and SNR for hybrid networks with exponential activations versus ReLU activations. Each baseline architecture is shown on a different row. Error bars represent the standard deviation of the mean across 5 random intializations.

## 4. Discussion

Here we demonstrate that 2 design principles for improving model interpretability for CNNs also generalizes to hybrid networks. Interestingly, we found that interpretability is largely driven by baseline module choices compared to the attention-based modules. We also found that networks that learn robust motif representations in first layer filters yield more reliable saliency maps. Learning distributed representations can result in noisier motifs which lead to learning a noisier function (Etmann et al., 2019). Such a function can maintain accurate predictions but may create noisy gradients, and hence less trustworthy attribution maps. Alternatively, learning better motifs may lead to learning a more robust and smoother function (Ilyas et al., 2019), thus providing more reliable gradients. In the future, we intend to expand this study to explore the interpretability of the attention maps and the role of positional encoding. Distance-dependent interactions are not relevant features for the synthetic data used in this study but can be important for regulatory function in real biological sequences. As attention-based models are gaining interest in regulatory genomics, hybrid networks would benefit from incorporating these design principles to bolster their intrinsic interpretability – first layer learns robust and identifiable motif representations while attention layers can focus on motif interactions – and increase their trustworthiness with gradient-based attribution methods.

